# *Trichoderma asperellum*: First report in Bangladesh

**DOI:** 10.1101/2021.08.13.456325

**Authors:** Sheikh Afsar Uddin, Ismail Hossain, Hyat Mahmud, M.S. Monjil, Mohammad Delwar Hossain

**Author notes:** **Corresponding author:** Hyat Mahmud.

## Abstract

The experiment was conducted at four laboratory, Bangladesh Agricultural University, Mymensingh to identify *T. asperellum* and its characterization was obtained from the rhizosphere of 49 different crops in 109 different locations of 36 districts in Bangladesh. Fifteen isolates of 88 isolates in *T. asperellum* were characterized on morphological and physiological features. All of the isolates were produced at 35 °C, but only TR_27_ and TR_45_ were grown and sporulate at 40 °C. Mycelial growth of all the isolates was reduced with the increasing of pH value. Molecular characterization in four isolates of TR_27_, TR_45_, TR_70_ and TR_85_ were studied at ITS and TEF region nucleotide sequencing among 15 isolates of *T. asperellum* and the morphological characterization at ITS and *tef1* nucleotide sequencing was assimilated with molecular characterization. The isolates of TR_45_, TR_70_ and TR_85_ were found with 98% homology and TR_27_ exhibited 88% against their respective closest isolate. The isolates of TR_27_ and TR_85_ marked their respective nearest homology at 96%, while TR_45_ showed 99%, and 93% homology was signified with TR_70_ in TEF sequences. Three isolates (TR_45_, TR_70_ and TR_85_) were identified as *T. asperellum* of 100% bootstrap value and TR_27_ isolate was also recognized with 72% bootstrap value in phylogenetic tree. In phylogenetic analysis, *tef1* extended better differentiation among the *Trichoderma* isolates within and among the groups of closely related species than the rDNA of the ITS region in reflecting wider variability in the isolates while a higher transversion ratio and evolutionary divergence were defined. Moreover the exploring of scientific and useful diversification of *Trichoderma* isolates on physiological and molecular traits has deemed in research fields.

**IMPORTANCE:** *Trichoderma* is an important genus in different aspect of uses as biofertilizer and bioremediation in agriculture. Specific identification in physiological and molecular characterization has also foremost attention with genome sequences as stored in databases. The diversity and genetic distances were ascertained among the isolates in phylogenetic analysis with the sequence data of the ITS region and *tef1* of known *Trichoderma* strains was found in GenBank. Consequently, the databases of *Trichoderma asperellum* in molecular characterization can be frequently studied in the beneficial scope of agriculture. *Trichoderma* isolates of Rhizosphere soil in 4 locations with crops of (TR_27_) Sadar, Moulvibazar (Rice), (TR_45_) Sadar, Mymensingh (Sweet gourd), (TR_70_) Chapra, Chapai Nawabganj (Sesame) and (TR_85_) Nayanpur, Lalmonirhat (Maize) were found to be substantiated as *Trichoderma asperellum* strain in Bangladesh that were confirmed on physiological and molecular characters, and utilization of this strain can run faster in invasive technologies to control crop diseases.

## INTRODUCTION

Identification of *Trichoderma* spp. is challenging in consequential to agricultural beneficial fungi. Pathogens are controlled by biological agents, because they are more natural and environmentally acceptable alternative to existing chemical treatments (1). Application of *Trichoderma* spp. in biofertilizers and biopesticides is used as bioeffectors (2, 3) and it also functions as bioremediation for heavy metals and other pollutions (4, 5).

*Trichoderma* has significant attributes as biocontrol agents (6) in controlling soil borne plant pathogenic fungi such as *Fusarium, Sclerotium* and *Rhizoctonia* (7, 8, 9). *Trichoderma* spp. also acts as an antagonistic effects over almost 18 genera and 29 species of pathogenic fungi, and a variety of pathogenic bacteria. *Trichoderma* spp. also in the biocontrol mechanisms exerts competition and mycoparasitism and often it causes the stimulation in plant resistance and immune system (10).

*Trichoderma* spp. shows mycoparasitism against pathogens in antagonistic mechanism. Firstly, *Trichoderma* recognizes the pathogen and its mycelium proceeds in straight way with pathogen mycelium in a coiling fashion that results in dissolution with death of the pathogen. As a result, cell wall-degrading enzymes (CWDEs), such as chitinases, glucanase, and proteases are produced due to the mycoparasitism, which penetrating the pathogen mycelium and nourishes its nutrients from dissolved pathogen (11, 12, 13). *Trichoderma* spp. also secrets secondary metabolites that induces the pathogen reduce. Different kind of substances as isonitrile, diketopiperazines, sesquiterpenes, stemids, polyketides, alkylpyrones, and peptaibols are produced from various *Trichoderma* species (14).

Several bioagents in 1980 were tested against plant pathogen but *Trichoderma* and, *Gliocladium* have gained the maximum success. So, bioagents have tremendous potential effect and can be exploited successfully in modern agriculture for control of plant diseases (15).

Effects of temperature on the growth and optimum sporulation of *Trichoderma viride and T. harzianum* are happened at 30-35 °C (16). Temperature ranges from 20 °C to 37 °C are favorable for best growth and sporulation, and growth and sporulation are also reduced at below 20 °C (17). Maximum growth and sporulation of *T. viride* are influenced between pH 4.5 to 5.5 (17). The highest growth and sporulation are also noticed at pH 5-7 (18). Potentiality of *Trichoderma* species as a bio-agent has been greatly impressed by pH, and the most favorable pH range of 6.5-7.5 is found in the growth and sporulation of *Trichoderma* (19).

Molecular characterization focuses on *Trichoderma* taxonomy and also in fungal diversification. In molecular study, eleven isolates of *T. viride* and eight isolates of *T. harzianum* are analyzed with ITS-PCR and ITS-PCR of rDNA region with ITS1 and ITS4 primers that produces 600 bp products in all isolates (20). *Trichoderma* isolates in different biocontrol capabilities characterize three main clades with ITS sequence analysis and reliable phylogenetic tree is constructed in the isolates of *T. harzianum* clade (21). Twelve isolates of *Trichoderma* spp. of different locations in South Andaman characterize with ITS-PCR and the sequence analysis of these isolates shows seven different species of *Trichoderma* spp. (22). Devi *et al*. (23) have defined the *Trichoderma* isolates as *T. virens* (11 isolates), *T. asperellum* (15), *T. harzianum* (14) and *T. longibrachiatum* (32) based on the DNA sequence data analysis of the elongation factor 1-alpha gene (*tef1*).

Significant effect of BAU-Biofungicide (*Trichoderma* based preparation) is observed in controlling the nursery diseases of litchi (24) and BAU-Biofungicide also exerts in inhibiting the leaf blight of wheat (25). BAU-Biofungicide is a unique biocontrol agent that greatly influence in higher germination and plant stand, low disease severity and enhancing grain yield of rice (26, 27).

The developments of efficient alternative measures are needed to combat diseases of crops for increasing yield. Although research on *Trichoderma spp*. has been carried out by the researchers in the country but detailed research on availability of biocontrol agents, their characterization and pathogenic study have not been carried out in different agro-ecological zones (AEZs). Our study reveals the resistance-related molecular mechanisms *of T. asperellum* with physiological and molecular characters. The present study has been designed to isolate, and identify with physiological and molecular characterization of *Trichoderma asperellum* collected from rhizosphere of 49 different crop species in 109 different locations of Bangladesh.

## RESULTS

### Selection of isolates for morphological and physiological characterization

Eighty eight isolates of *Trichoderma* were grouped in 49 crops with 109 locations of the country. The isolates were categorized as fast, medium and slow growing on the basis of their growth habit. Fifteen isolates were selected from the fast growing *Trichoderma* isolates on the basis of different rhizosphere crops in different locations (Table 1).

**Table 01.** Spore density of different isolates of *Trichoderma* at 30°C.

| Isolate of *Trichoderma* | Location | Name of crop & scientific name | No. of spore/ml |  |  |
| --- | --- | --- | --- | --- | --- |
|  |  |  | 7 DOC | 14 DOC | 21 DOC |
| TR<sub>9</sub> | Dashmina, Patuakhali | Brinjal (*Solanum melongena*) | 4.74×10<sup>7</sup>cd | 5.83×10<sup>7</sup>de | 7.50×10<sup>7</sup>c |
| TR<sub>11</sub> | Kawkhali, Pirojpur | Taro (*Colocasia esculenta*) | 4.65×10<sup>7</sup>d | 5.58×10<sup>7</sup>ef | 5.71×10<sup>7</sup>c |
| TR<sub>25</sub> | Zaflang, Sylhet | Mustard (*Brassica nigra*) | 4.55×10<sup>7</sup>d | 5.43×10<sup>7</sup>ef | 5.79e×10<sup>7</sup>f |
| TR<sub>27</sub> | Sadar, Moulvibazar | Rice (*Oryza sativa*) | 5.81×10<sup>7</sup>a | 7.35×10<sup>7</sup>b | 9.55×10<sup>7</sup>a |
| TR<sub>45</sub> | Sadar, Mymensingh | Sweet gourd (*Cucurbita muschata*) | 5.61×10<sup>7</sup>ab | 7.44×10<sup>7</sup>b | 9.37×10<sup>7</sup>ab |
| TR<sub>52</sub> | Sadar, Mymensingh | Papaya (*Carica papaya*) | 4.86×10<sup>7</sup>cd | 6.43×10<sup>7</sup>c | 6.33×10<sup>7</sup>ef |
| TR<sub>56</sub> | Melando, Jamalpur | Brinjal (*Solanum melongena*) | 4.38×10<sup>7</sup>d | 4.54×10<sup>7</sup>g | 7.15×10<sup>7</sup>c |
| TR<sub>64</sub> | Shipganj, Bogra | Maize (*Zea mays*) | 4.91×10<sup>7</sup>cd | 6.45×10<sup>7</sup>c | 7.25×10<sup>7</sup>c |
| TR<sub>70</sub> | Chapra, Chapai nawabganj | Sesame (*Sesamum indicum*) | 5.92×10<sup>7</sup>a | 7.56×10<sup>7</sup>b | 8.82×10<sup>7</sup>b |
| TR<sub>77</sub> | Sadar, Rangpur | Rice (*Oryza sativa*) | 4.99×10<sup>7</sup>cd | 5.59×10<sup>7</sup>ef | 6.05×10<sup>7</sup>ef |
| TR<sub>78</sub> | Gobindoganj, Gaibanda | Tobacco (*Nicotiana tabacum*) | 4.68×10<sup>7</sup>cd | 5.24×10<sup>7</sup>f | 6.30×10<sup>7</sup>ef |
| TR<sub>79</sub> | Sadar, Kurigram | Wheat (*Triticum aestivum*) | 5.34×10<sup>7</sup>bc | 6.30×10<sup>7</sup>cd | 7.47×10<sup>7</sup>c |
| TR<sub>82</sub> | Sadar, Kurigram | Okra (*Abelmoschus esculentus*) | 4.39×10<sup>7</sup>d | 5.79×10<sup>7</sup>de | 7.08×10<sup>7</sup>cd |
| TR<sub>85</sub> | Nayanpur, Lalmonirhat | Maize (*Zea mays*) | 5.85×10<sup>7</sup>a | 8.66×10<sup>7</sup>a | 9.05×10<sup>7</sup>ab |
| TR<sub>87</sub> | Pirganj, Thakargaon | Natishak *Brassica chinensis* | 4.90 ×10<sup>7</sup> cd | 5.56×10<sup>7</sup>ef | 6.43×10<sup>7</sup>ed |
| CV (%) |  |  | 8.12 | 5.47 | 5.37 |

Data in column having same letter(s) do not differ significantly at 1% level of significance; DOC = Days old culture

### Spore density of different isolates of *Trichoderma* at 30 °C

Fifteen fast growing isolates of *Trichoderma* were observed for the number of spore per ml soil at 30 °C temperature in 3 times such as 7, 14 and 21 days old culture (DOC). The number of spore per ml was increased with the increasing of the age of the culture. Significant variation was found among the spore density of the isolates. The highest number of spores was obtained in the isolates of maize field of Nayanpur upazilla, Lalmonirhat district at 14 and 21 days old culture. Another higher number of spores was recorded at 21 DOC in T_27_ that was collected from rice field of Sadar upazilla, Moulvibazar district (Table 1).

### Effect of temperature on radial mycelial growth of different *Trichoderma* isolates

Radial mycelial growth and spore density of *Trichoderma* were increased as the increasing of temperatures, but spore density was decreased at 35 °C (Table 3). Four isolates such as Sadar, Moulvibazar,, Sadar, Mymensingh, Chapra, Chapai nawabganj and Nayanpur, Lalmonirhat produced higher mycelial growth and spore density on PDA at different temperatures and different ages of the culture. Only two (TR_27_ and TR_45_) isolates of *Trichoderma* were grown at 40 °C which were collected from Sadar upazilla, Moulvibazar district and Sadar Mymensingh, respectively (Fig. 01). The 15 isolates of *Trichoderma* prolonged regular shaped colony under different temperatures viz. 15 °C, 20 °C, 25 °C and 30 °C. All of the *Trichoderma* isolates showed five different colony colour viz. whitish, whitish green, green, dark green and light green and two types of colony consistency such as compact and loose were observed at different temperatures (Table 2).

**Fig.01.**
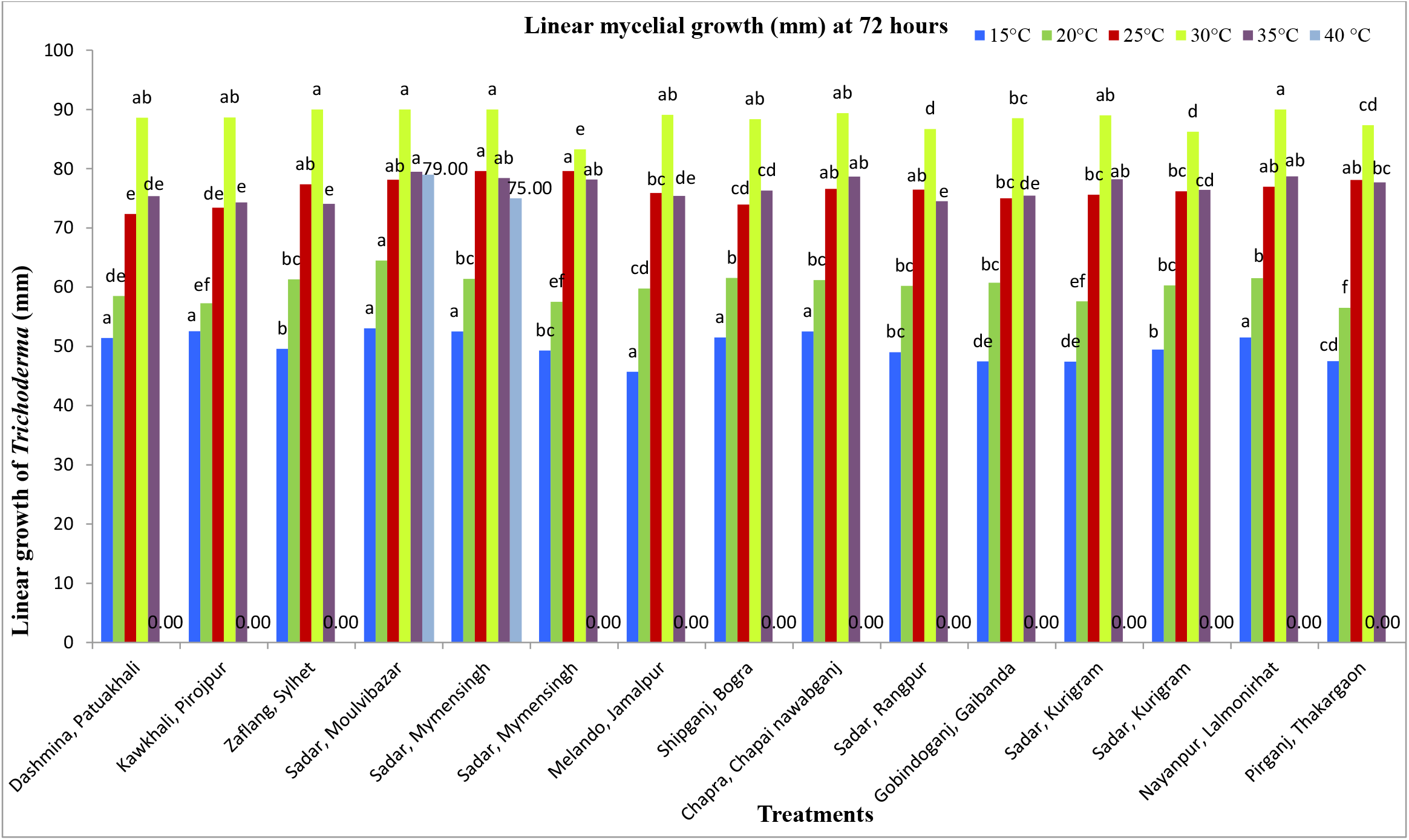
Effect of different temperatures on the radial mycelial growth of different isolates of *Trichoderma* in PDA at 72 hours.

**Table 02.** Effect of different temperatures on colony growth characters of different *Trichoderma* isolates on PDA after 72 hrs.

| Isolate of<br>*Trichoderma* | Colony color |  |  |  |  |  | Colony compactness |  |  |  |  |
| --- | --- | --- | --- | --- | --- | --- | --- | --- | --- | --- | --- |
|  | 15°C | 20°C | 25°C | 30°C | 35°C | 40°C | 15°C | 20°C | 25°C | 30°C | 35°C |
| TR<sub>9</sub> | W | W | WG | G | G | - | L | L | L | C | C |
| TR<sub>11</sub> | W | G | G | G | DG | DG | L | L | C | C | C |
| TR<sub>25</sub> | W | WG | WG | WG | WG | WG | L | C | C | L | L |
| TR<sub>27</sub> | W | W | WG | G | G | - | L | L | C | C | C |
| TR<sub>45</sub> | W | WG | WG | G | G | - | L | L | C | C | C |
| TR<sub>52</sub> | W | W | WG | G | G | - | L | L | L | C | C |
| TR<sub>56</sub> | W | WG | WG | G | G | - | L | L | L | C | C |
| TR<sub>64</sub> | W | W | WG | WG | LG | - | L | C | C | L | L |
| TR<sub>70</sub> | W | WG | WG | G | G | - | L | L | L | C | C |
| TR<sub>77</sub> | W | W | WG | G | G | - | L | L | L | L | L |
| TR<sub>78</sub> | W | W | WG | G | G | - | L | L | C | C | L |
| TR<sub>79</sub> | W | W | WG | WG | LG | - | L | L | C | L | C |
| TR<sub>82</sub> | W | W | WG | G | WG | - | L | L | L | L | L |
| TR<sub>85</sub> | W | W | WG | G | DG | - | L | L | C | C | C |
| TR<sub>87</sub> | W | W | WG | WG | G | - | L | L | C | L | C |

**W** = Whitish, **WG** = Whitish Green, **G** = Green, **DG** = Dark Green, **LG** = Light Green,
**L** = Loose, **C** = Compact; - = no growth of *Trichoderma*

**Table 03.** Spore density of different *Trichoderma* on PDA at different temperatures of 7 days old culture.

| Isolate of<br>*Trichoderma* | Location | Crop | Contain of spore/ml |  |  |
| --- | --- | --- | --- | --- | --- |
|  |  |  | 25<sup>0</sup>C | 30<sup>0</sup>C | 35<sup>0</sup>C |
| TR<sub>9</sub> | Dashmina, Patuakhali | Brinjal | 4.95×10<sup>7</sup>cd | 5.61×10<sup>7</sup>d | 2.83×10<sup>7</sup>e |
| TR<sub>11</sub> | Kawkhali, Pirojpur | Taro | 4.59×10<sup>7</sup>de | 5.81×10<sup>7</sup>d | 2.92×10<sup>7</sup>e |
| TR<sub>25</sub> | Zaflang, Sylhet | Mustard | 4.40×10<sup>7</sup>e | 6.85×10<sup>7</sup>bc | 3.29×10<sup>7</sup>de |
| TR<sub>27</sub> | Sadar, Moulvibazar | Rice | 5.91×10<sup>7</sup>b | 7.36×10<sup>7</sup>ab | 5.57×10<sup>7</sup>a |
| TR<sub>45</sub> | Sadar, Mymensingh | Sweet gourd | 7.05×10<sup>7</sup>a | 7.85×10<sup>7</sup>a | 5.63×10<sup>7</sup>a |
| TR<sub>52</sub> | Sadar, Mymensingh | Papaya | 4.99×10<sup>7</sup>cd | 6.62×10<sup>7</sup>c | 4.62×10<sup>7</sup>b |
| TR<sub>56</sub> | Melando, Jamalpur | Brinjal | 4.63×10<sup>7</sup>de | 5.26×10<sup>7</sup>de | 4.80×10<sup>7</sup>b |
| TR<sub>64</sub> | Shipganj, Bogra | Maize | 4.58×10<sup>7</sup>de | 4.60×10<sup>7</sup>e | 3.53×10<sup>7</sup>cd |
| TR<sub>70</sub> | Chapra, Chapai<br>Nababganj | Sesame | 5.17×10<sup>7</sup>c | 7.65×10<sup>7</sup>a | 4.78×10<sup>7</sup>b |
| TR<sub>77</sub> | Sadar, Rangpur | Rice | 4.71×10<sup>7</sup>cd | 5.79×10<sup>7</sup>d | 4.58×10<sup>7</sup>b |
| TR<sub>78</sub> | Gobindoganj, Gaibanda | Tobacco | 5.16×10<sup>7</sup>c | 5.55×10<sup>7</sup>d | 3.88×10<sup>7</sup>c |
| TR<sub>79</sub> | Sadar, Kurigram | Wheat | 4.65×10<sup>7</sup>de | 6.63×10<sup>7</sup>c | 3.75×10<sup>7</sup>cd |
| TR<sub>82</sub> | Sadar, Kurigram | Okra | 4.37×10<sup>7</sup>e | 6.68×10<sup>7</sup>c | 4.76×10<sup>7</sup>b |
| TR<sub>85</sub> | Nayanpur, Lalmonirhat | Maize | 4.95×10<sup>7</sup>cd | 7.69×10<sup>7</sup>a | 5.07×10<sup>7</sup>ab |
| TR<sub>87</sub> | Pirganj, Thakargaon | Natishak | 4.80×10<sup>7</sup>cd | 6.91×10<sup>7</sup>bc | 4.77×10<sup>7</sup>b |
| CV (%) |  |  | 5.54 | 6.17 | 8.13 |

Data in column having same letter(s) do not differ significantly at 1% level of significance.

### Influence of pH on mycelia of different isolates of *Trichoderma*

Four different pH viz., 6.0, 7.0, 8.0 and 8.5 were selected to study the influence of pH on the mycelial growth on PDA among 15 isolates of *Trichoderma*. Fifteen isolates of *Trichoderma* showed higher average radial mycelial growth rate at pH 6.0 and pH 7.0 among the tested pH. The radial mycelial growth rate was reduced with the higher value of pH at 8.00 and 8.5. Four isolates such as Sadar Moulvibazar, Sadar Mymensingh, Chapra, Chapai nawabganj and Nayanpur, Lalmonirhat were found to have better mycelial growth on PDA at different pH values in the isolates of *Trichoderma* (Fig. 2). The 15 isolates of *Trichoderma* produced regular and irregular shaped colony in different p^H^ values at 6.0, 7.0, 8.0 and 8.50. Four different colony colours viz. whitish, green, dark green and light green were observed in the fifteen isolates. Two types of colony consistency were obtained such as compact and loose at different p^H^ values (Table 4).

**Fig. 02.**
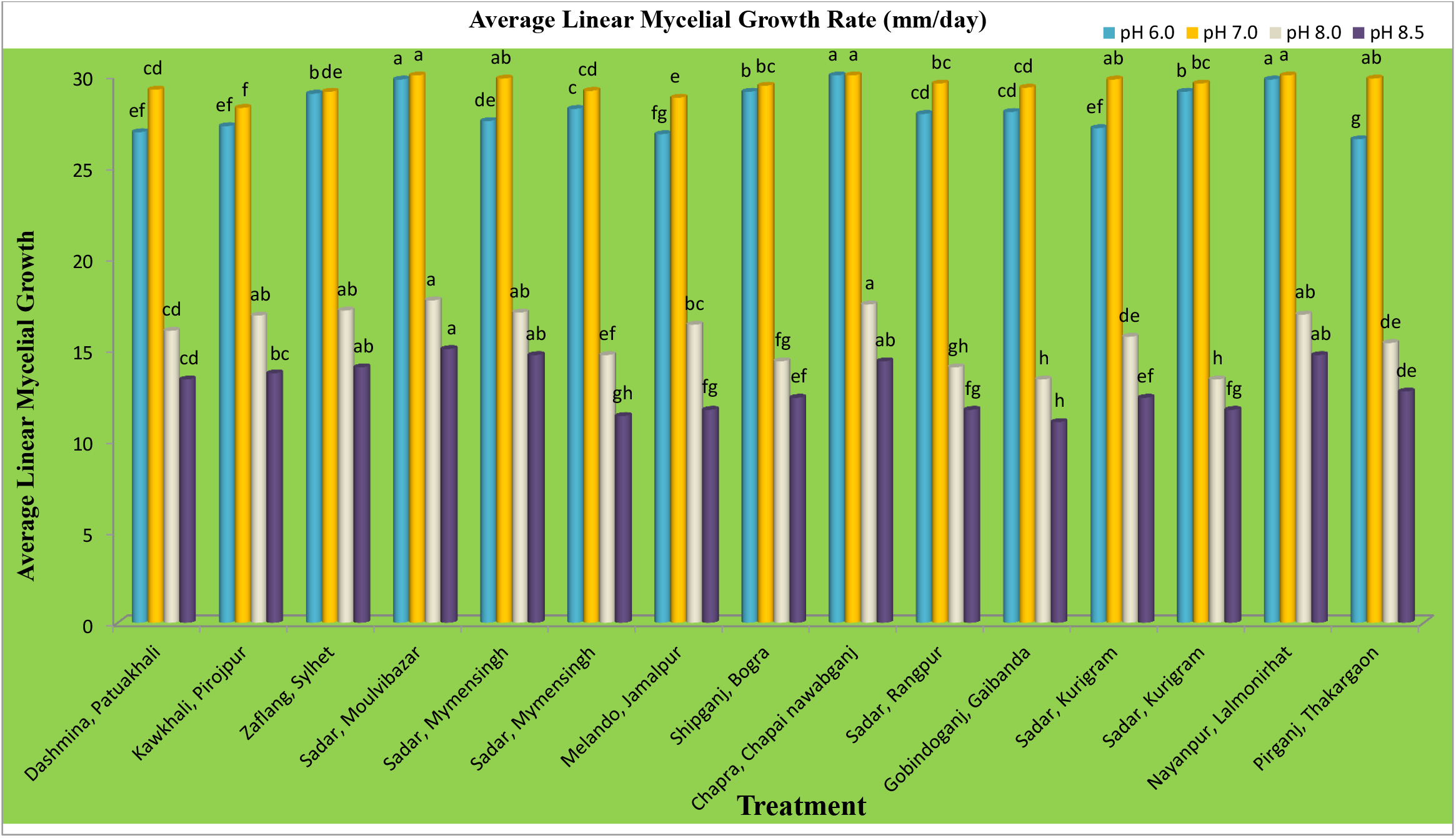
Effect of pH on the growth of different isolates of *Trichoderma*.

**Table 04.** Effect of different pH levels oncolony growth characters of different isolates of *Trichoderma* on PDA.

| Isolate of *Trichoderma* | Colony growth characters at different p<sup>H</sup> level |  |  |  |  |  |  |  |  |  |  |  |
| --- | --- | --- | --- | --- | --- | --- | --- | --- | --- | --- | --- | --- |
|  | Colony shape |  |  |  | Colony color |  |  |  | Colony compactness |  |  |  |
|  | p<sup>H</sup> =6 | p<sup>H</sup> =7 | p<sup>H</sup> =8 | p<sup>H</sup> =8.5 | p<sup>H</sup> =6 | p<sup>H</sup> =7 | p<sup>H</sup> =8 | p<sup>H</sup> =8.5 | p<sup>H</sup> =6 | p<sup>H</sup> =7 | p<sup>H</sup> =8 | p<sup>H</sup> =8.5 |
| TR<sub>9</sub> | Regular | Regular | Regular | Irregular | Green | Green | Green | Green | Compact | Compact | Compact | Compact |
| TR<sub>11</sub> | Regular | Regular | Regular | Irregular | Whitish | Green | Whitish | Whitish | Loose | Compact | Compact | Compact |
| TR<sub>25</sub> | Irregular | Regular | Regular | Regular | Green | Green | Green | Green | Loose | Compact | Compact | Compact |
| TR<sub>27</sub> | Regular | Regular | Regular | Irregular | Green | Green | Green | Green | Compact | Compact | Compact | Compact |
| TR<sub>45</sub> | Regular | Regular | Regular | Regular | Green | Green | Green | Green | Compact | Compact | Compact | Compact |
| TR<sub>52</sub> | Regular | Regular | Regular | Regular | Green | Green | Whitish | Green | Compact | Compact | Compact | Compact |
| TR<sub>56</sub> | Irregular | Regular | Irregular | Irregular | Whitish | Green | Whitish | Green | Loose | Compact | Loose | Loose |
| TR<sub>64</sub> | Irregular | Regular | Irregular | Irregular | Light green | Green | Light green | Light green | Compact | Compact | Compact | Compact |
| TR<sub>70</sub> | Regular | Regular | Regular | Regular | Green | Green | Green | Green | Compact | Compact | Compact | Compact |
| TR<sub>77</sub> | Regular | Regular | Regular | Regular | Green | Green | Green | Green | Compact | Compact | Compact | Compact |
| TR<sub>78</sub> | Regular | Regular | Regular | Regular | Green | Green | Green | Green | Compact | Compact | Compact | Compact |
| TR<sub>79</sub> | Irregular | Regular | Irregular | Irregular | Light green | Green | Green | Light green | Compact | Compact | Compact | Compact |
| TR<sub>82</sub> | Regular | Regular | Regular | Regular | Green | Green | Green | Green | Compact | Compact | Compact | Compact |
| TR<sub>85</sub> | Regular | Regular | Regular | Regular | Dark green | Green | Green | Green | Compact | Compact | Compact | Loose |
| TR<sub>87</sub> | Regular | Regular | Regular | Regular | Green | Green | Green | Green | Compact | Compact | Compact | Compact |

### Molecular features of selected *Trichoderma* isolates

Four (TR_27,_ TR_45_,TR_70_ and TR_85_) *Trichoderma* Isolates were further studied for molecular characterization on the basis of their mycelial growth, morphological and physiological characters at tolerable levels to high temperature and high spore density per ml. These four isolates of *Trichoderma* were identified on the basis of their spore and mycelial structure (Fig.03.) and characterized at molecular level on the basis of PCR amplification at ITS4, ITS5, *tef1* and *tef2* region.

**Fig. 03.**
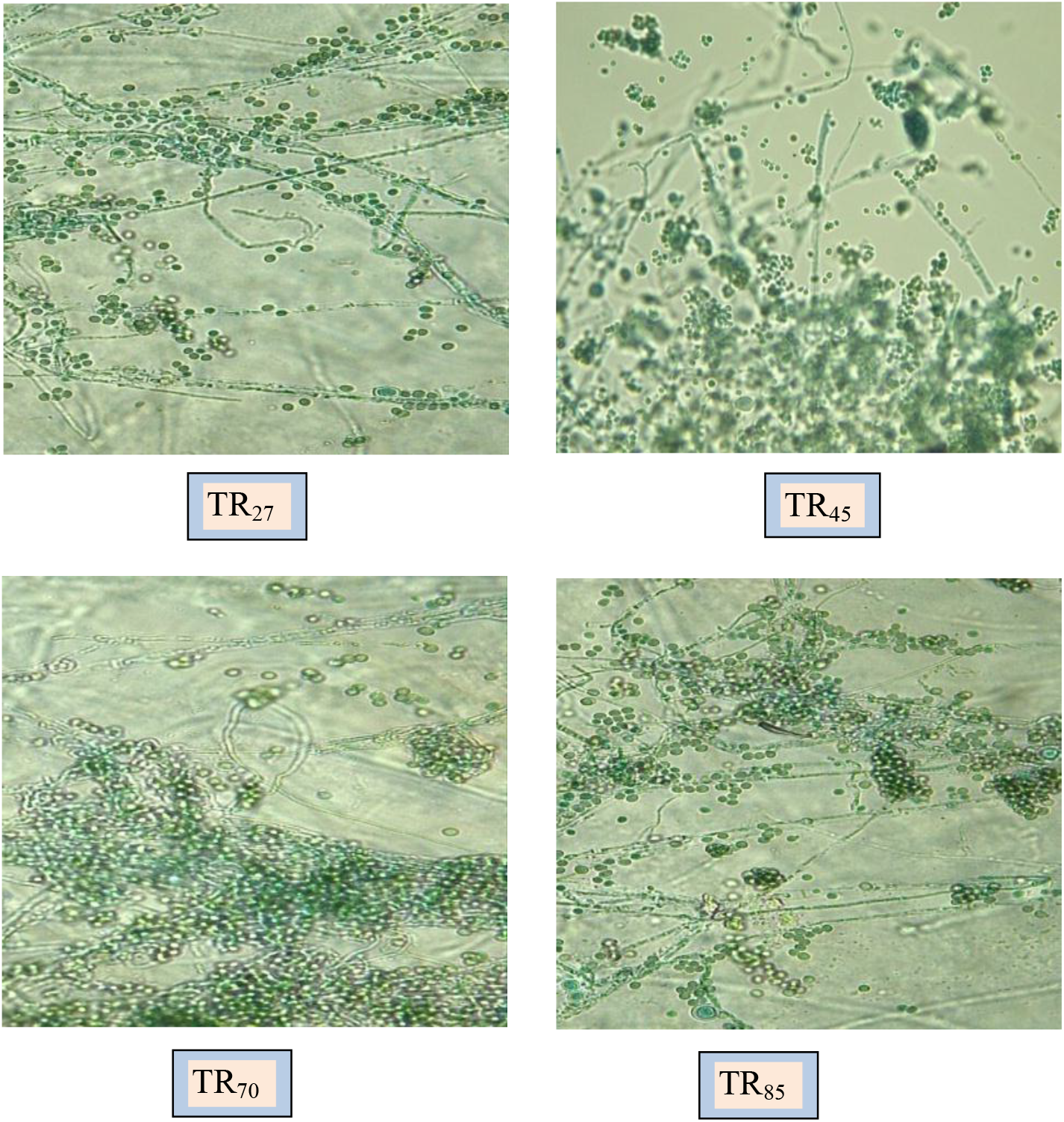
Structure of different isolates of *Trichoderma asperellum*.

### Molecular characterization

#### Characterization based on ITS region

Sequence analyses of ITS region in nucleotide PCR amplificaion at ITS4 region and ITS5 region resulted in 650 bp size band (Fig.04 A). Blast homology revealed that the isolate of Sadar, Moulvibazar showed 88% homology with *Trichoderma asperellum* India (Accession number : KY-401447.1), while Chapra, Chapai nawabganj pronounced 98% homology on *Trichoderma asperellum* India (Accession number: KT-001078.1). Ninety eight percent homology of *Trichoderma asperellum* Mexico (Accession number: KP-340278.1) was observed in the isolate of Sadar, Mymensingh and the isolate of Nayanpur, Lalmonirhat exhibited 98% homology with *Trichoderma asperellum* Mexico (Accession number: KP-340257.1) as presented in Table 5. Four isolates were determined as *T. asperellum* in the molecular sequence at ITS region from the above findings.

**Fig. 04.**
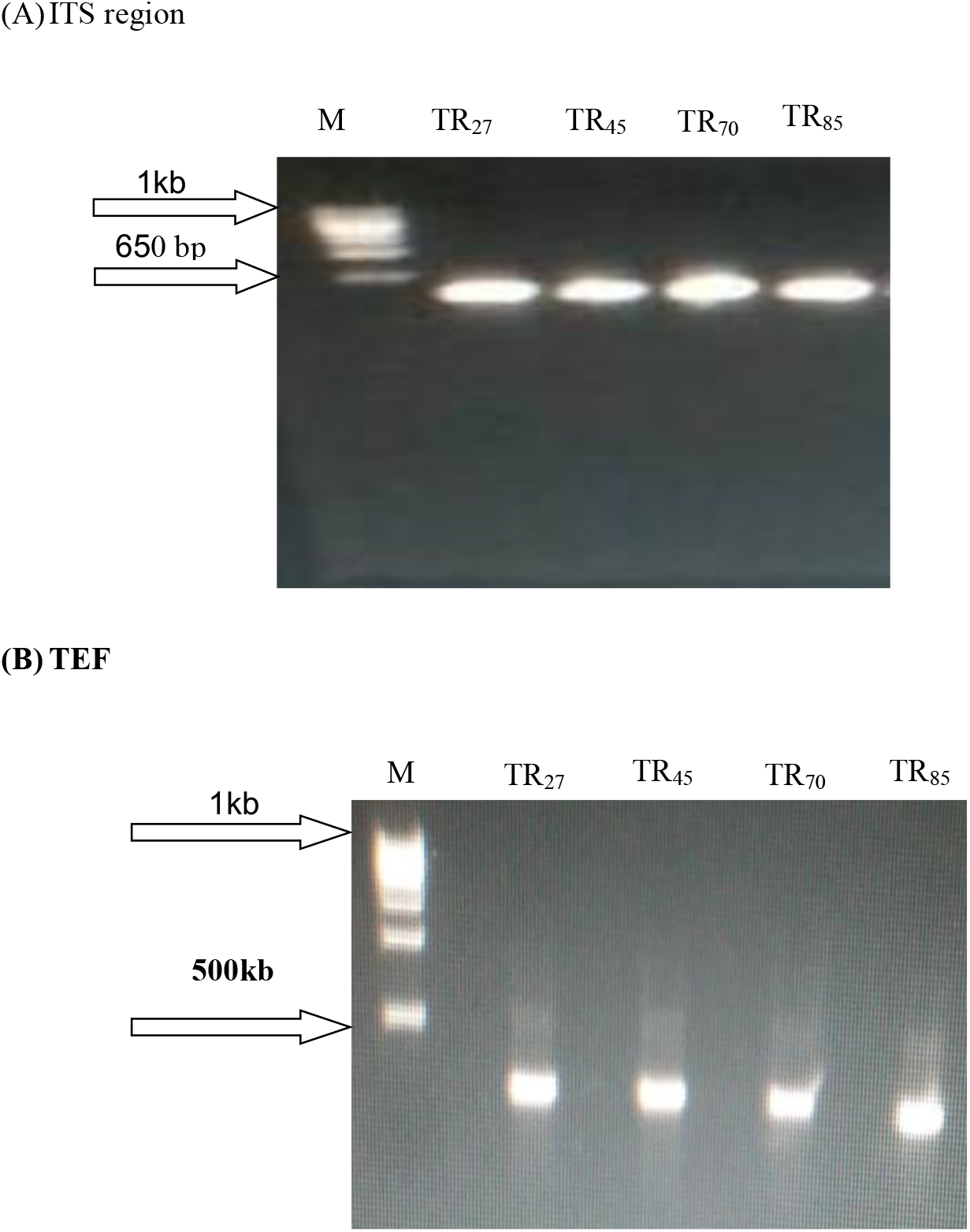
PCR amplification of ITS region of *Trichoderma* isolates using ITS4 and ITS5 primers (A) ITS region and the *tef1* and *tef2* primers for TEF (B). Lanes: TR_27_, TR_45_, TR_70_ and TR_85_ isolates, respectively. M: DNA ladder size. TR_27_ : Moulvi Bazar, TR_45_ : Mymensingh, TR_70_ : Chapainababganj and TR_85_ : Lalmonirhat.

**Fig. 05.**
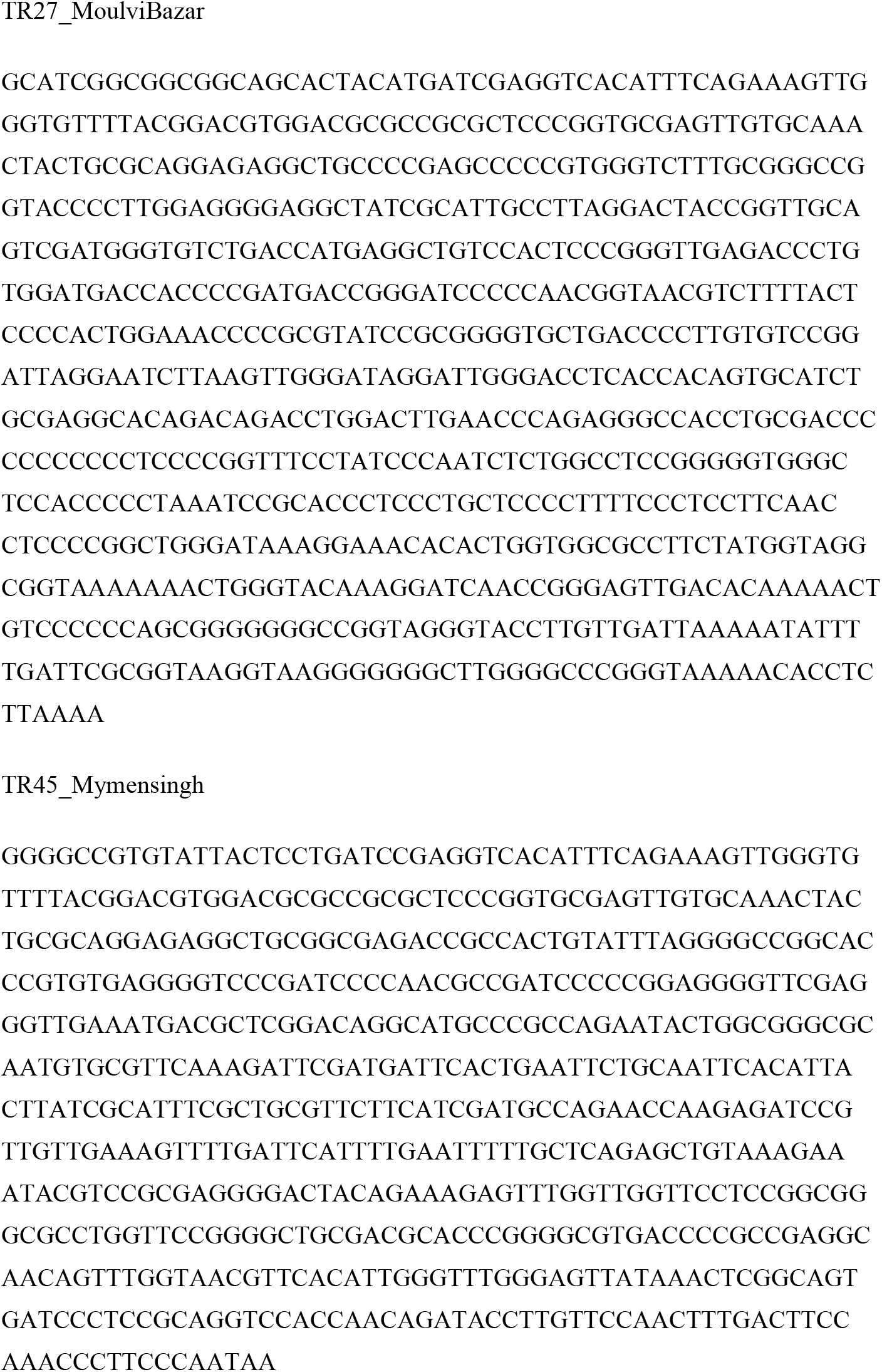
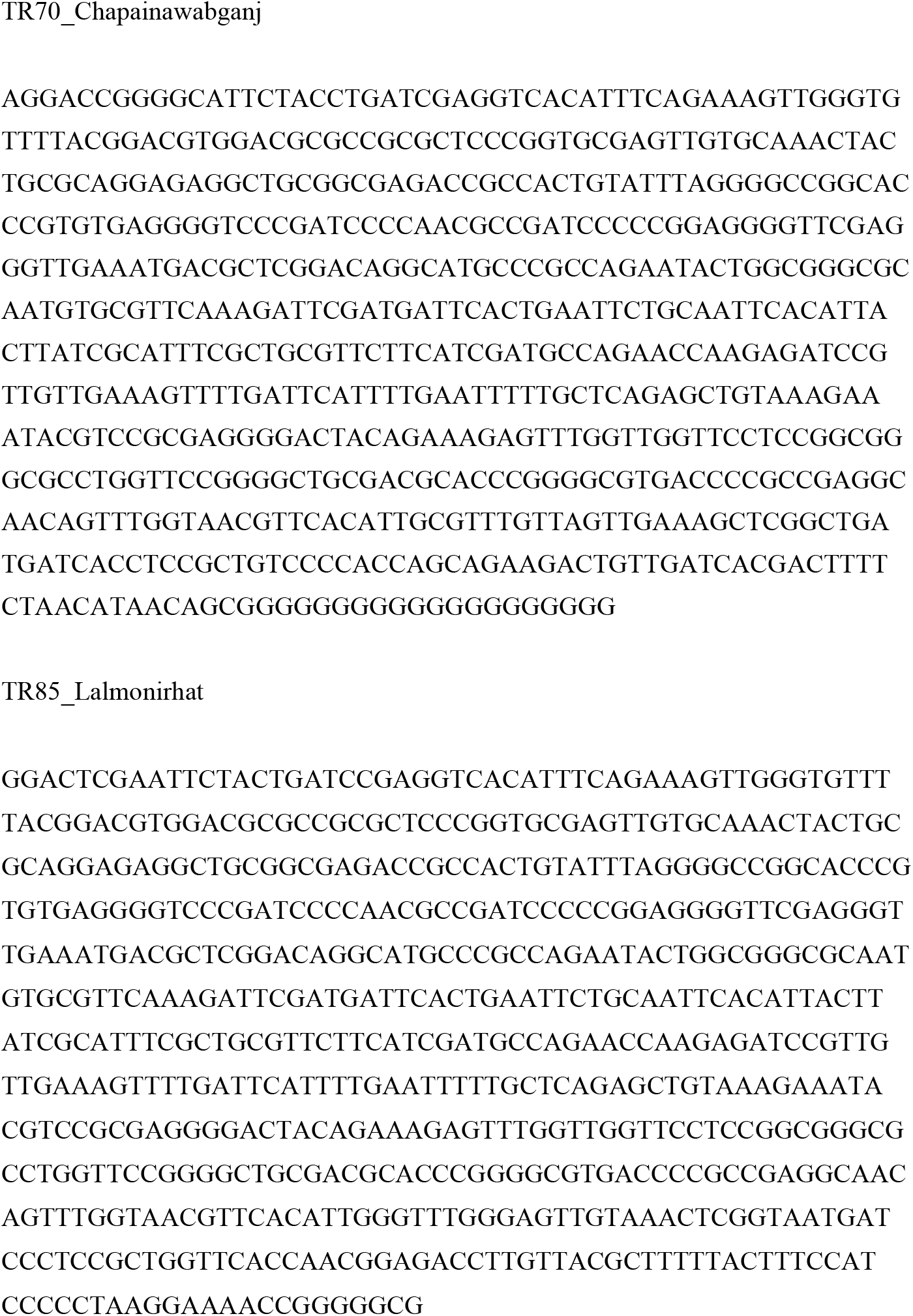
Sequence of ITS region of different isolates of *Trichoderma asperellum*.

**Fig. 06.**
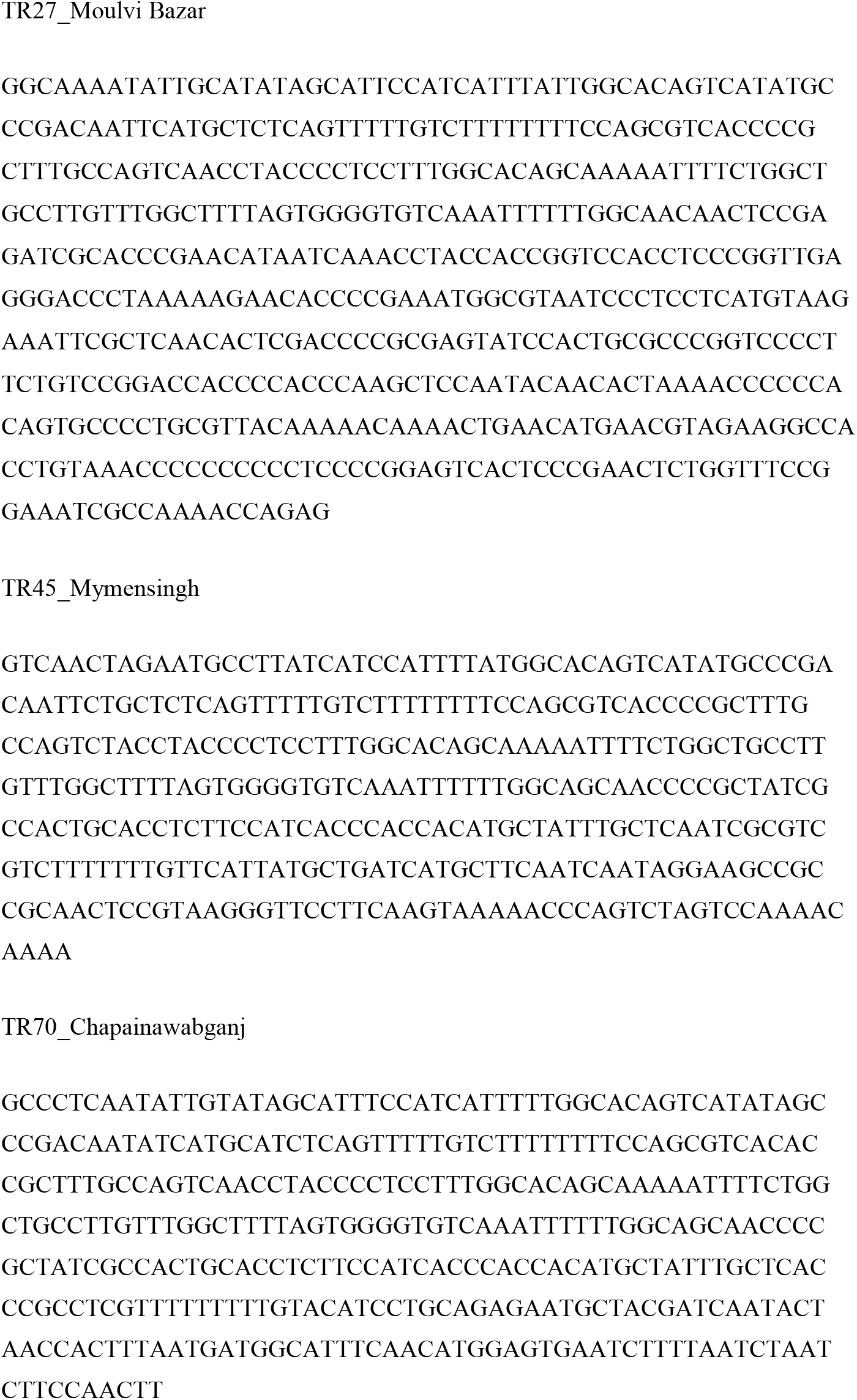
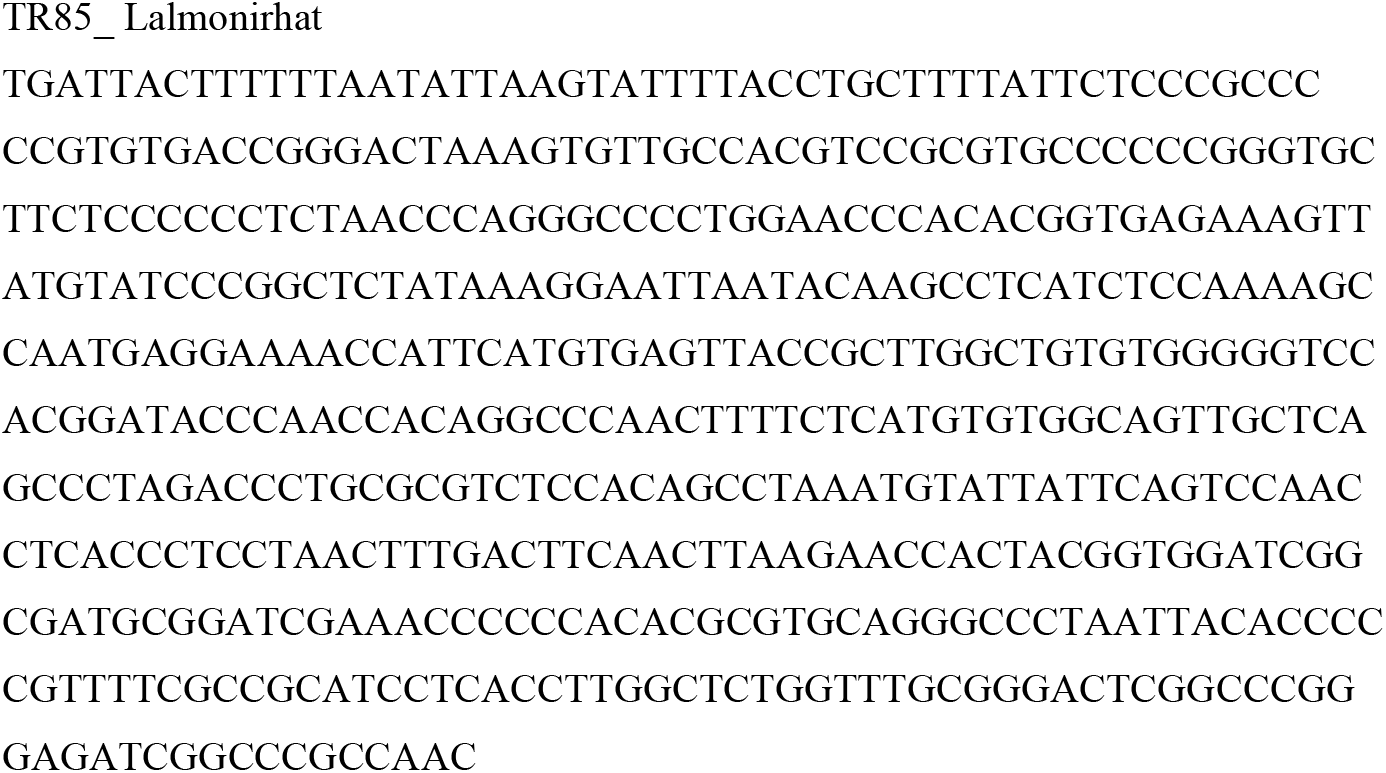
Sequence of TEF region of different isolates of *Trichoderma asperellum*.

**Table 5.**
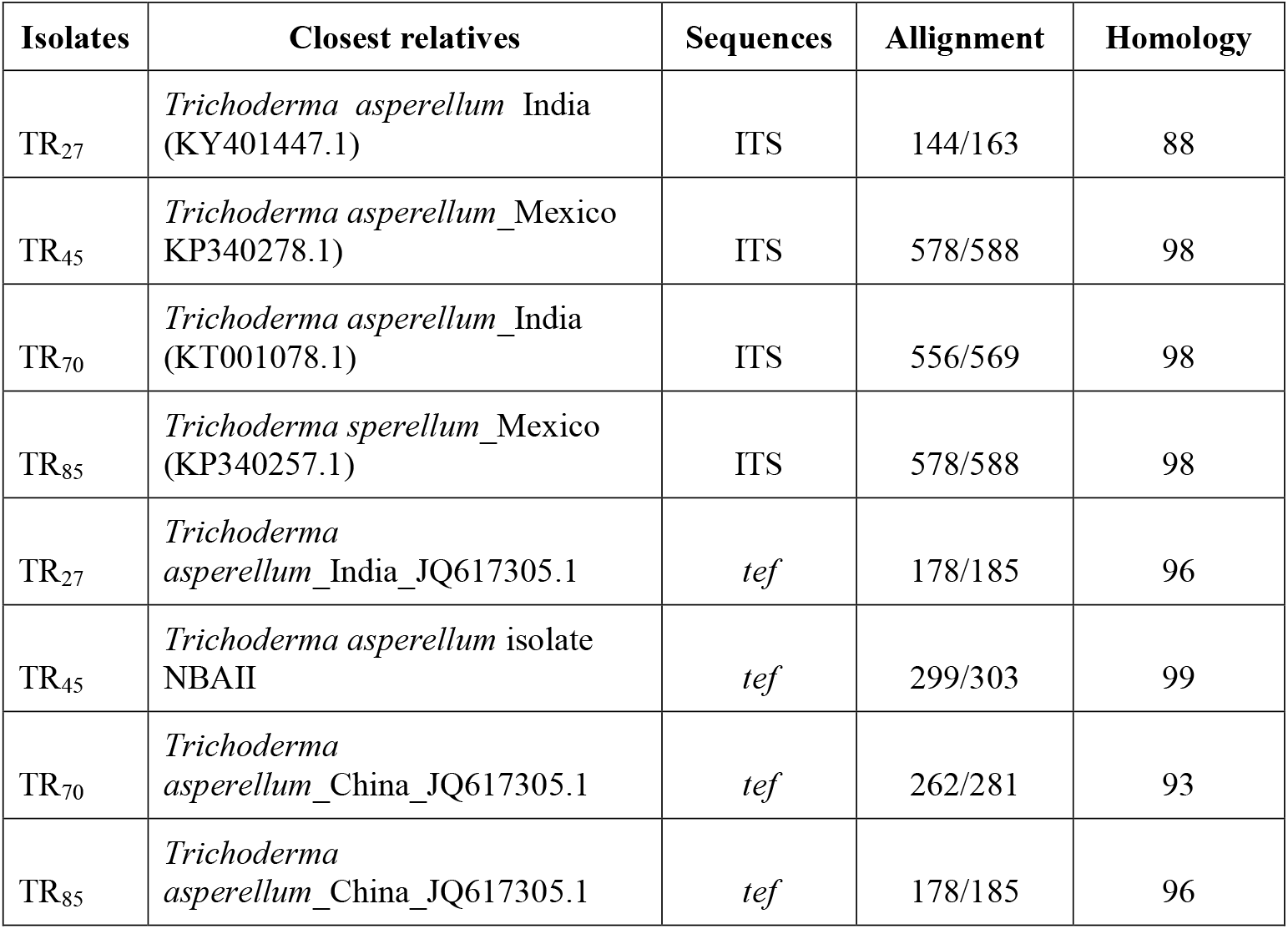
Closest relatives of the selected *Trichoderma* isolates based on ITS and *tef* sequences.

### Characterization based on *TEF*

The *Trichoderma* isolates were also identified by sequencing of *tef1* gene. PCR amplification with *tef*1 and *tef*2 region showed 350 bp band (Fig. 04 B). Isolates of Sadar, Moulvibazar, Chapra, Chapai nawabganj, Sadar, Mymensingh and Nayanpur, Lalmonirhat were defined as *T. asperellum* based on the sequence analyses of *tef*1 in Blast program. The isolate of Sadar, Moulvibazar appeared at 96% homology with *Trichoderma asperellum* India (Accession number: JQ-617305.1), *Trichoderma asperellum* NBAII (Accession number: GU-592425.1) was found to be 99% homology in the isolate of Chapra, Chapai nawabganj. Sadar, Mymensingh marked 93% homology on *Trichoderma asperellum* China (Accession number: JQ-617305.1) and 96% homology was found with *Trichoderma asperellum* China (Accession number: JQ-617305.1) in Nayanpur, Lalmonirhat isolate (Table 5).

### Phylogenetic tree of selected *Trichoderma* based on ITS and *tef* sequences

The phylogenetic tree differentiated the four *Trichoderma* isolates based on the ITS sequences in two clusters (Fig. 07). Cluster I consisted of 3 isolates such as Sadar, Mymensingh, Chapra, Chapai nawabganj and Nayanpur, Lalmonirhat were ascertained as *T. asperellum* that was supported by a bootstrap value of 100% and the isolate of Sadar, Moulvibazar isolate (Sub cluster II) was also noticed as *T. asperellum* and it was supported in bootstrap of 72%. Phylogenetic tree in the *Trichoderma asperellum* was also divided into two clusters at *tef* sequences whilst cluster I was grouped into two sub clusters. In cluster I, the isolate of Sadar, Mymensingh was supported by 98% bootstrap value and 99% bootstrap value was found in the isolate of Chapra, Chapai nawabganj. In sub-cluster II, the isolate in Moulvibazar district of Bangladesh was observed, whereas the isolate of Nayanpur, Lalmonirhat was grouped in cluster II with 100% bootstrap value (Fig. 08).

**Fig. 07.**
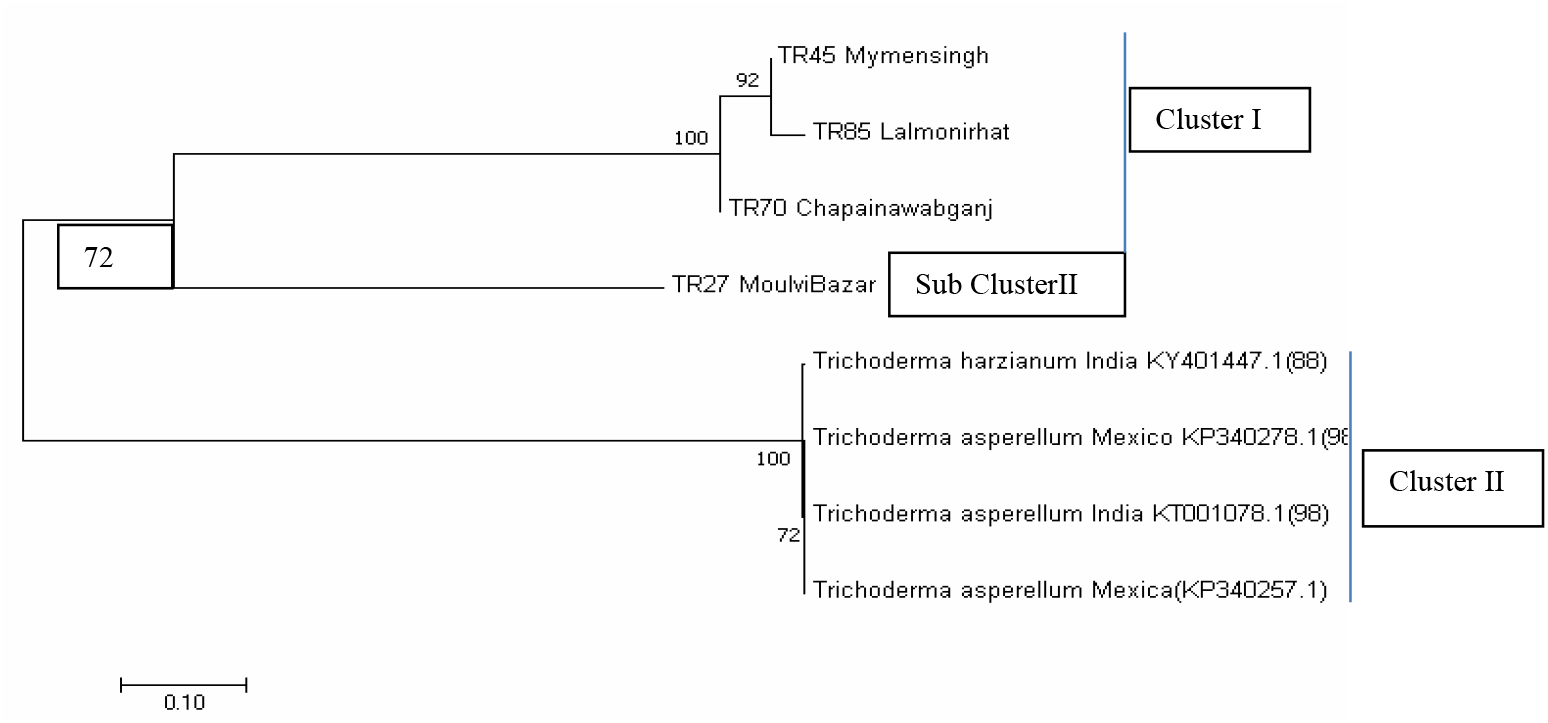
Phylogenetic relationships among the *Trichoderma* isolates in the analysis of ITS sequences.

**Fig. 08.**
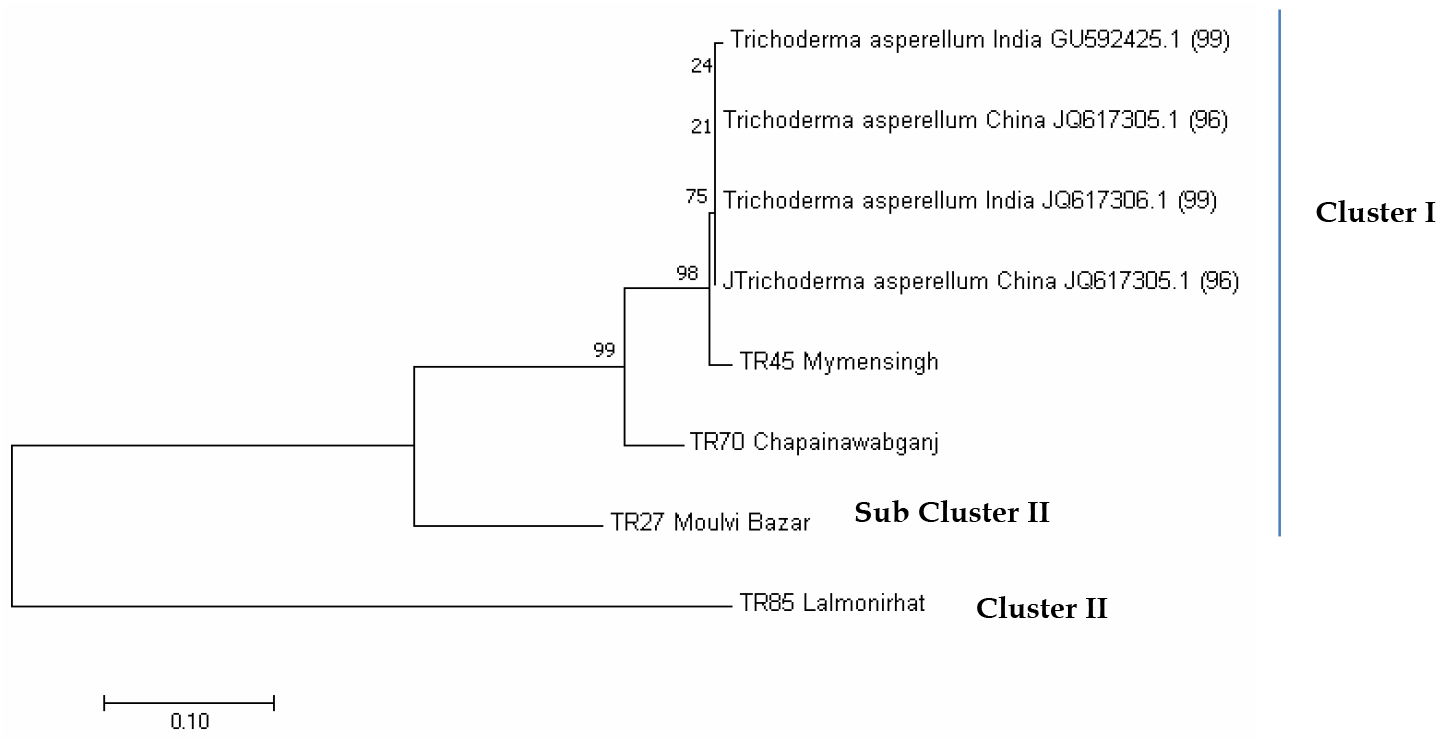
Phylogenetic relationships among the *Trichoderma* isolates by analysis of *tef* sequences.

## DISCUSSION

### Analysis and evaluation of spore density of different isolates of *Trichoderma* at 30 °C

Eighty eight isolates of *Trichoderma* were collected from rhizosphere soils of 49 different crops 109 different locations of thirty six districts in Bangladesh. Colony color, radial growth, growth habit, consistency, conidia and spore density were found to show distinct variable. These findings were supported by the researcher (28) who reported that morphological characteristics showed significant variation among different isolates. They differed in mycelial growth, colony color, colony consistency and sporulation rate (29). The morphological differences when grown on PDA may be related to nutritional factors, including environmental and genetic factors influencing fungal growth and development. Fungal growth was affected over a large environmental range (30) where BCAs were sensitive to environmental conditions that had been cited as a reason for inconsistent performance (31).

### Effect of temperature on radial mycelial growth of different *Trichoderma* isolates

We found that temperature and pH greatly influenced the mycelial growth and sporulation of the *Trichoderma* isolates. All of the *Trichoderma* isolates revealed the highest radial mycelial growth at 30 °C temperature followed by 15 °C, 20 °C, 25 °C and 35 °C. Similar results were supported by the scientists (17, 32, 33). The most favorable temperature for the growth of *Trichoderma* species was between 25 - 30 °C (34). Growth and sporulation of *Trichoderma harzianum* was found to be at 30 °C followed by 25° C (19). The fungal colony growth of antagonistic microbes *T. harzianum* and *T. pseudokongii* on PDA was determined at optimum 30 °C (35).

### Evaluation and influence of pH on mycelia of different isolates of *Trichoderma* isolates

The highest radial mycelial growth was attained at p^H^ 6.0 and p^H^ 7.0 among all the isolates of *Trichoderma*. This result was in agreement with the findings (36, 37), who reported that similar results were observed with different *Trichoderma* isolates. The researchers (33) also reported that the most favorable pH range for *Trichoderma* was between 5.5 and 7.5. The best growth of *Trichoderma virens* was noted at pH 5.5 and optimal growth was between 5.0 and 7.0 (38). Potentiality of *Trichoderma* species as a bioagent that has been greatly influenced by pH and most favorable pH range was 6.5-7.5 for growth and sporulation (19).

### Analytical assessment on molecular features of *Trichoderma asperellum*

*Trichoderma* isolates were proved by sequencing of ITS and TEF gene region in this study. The molecular characteristics of the ITS region and tef1gene of the *Trichoderma* isolates revealed that the four *Trichoderma* isolates were divided in two clusters. Similar findings were observed by the reporter (22) who observed 12 isolates of *Trichoderma* in different locations of South Andaman with ITS region that were characterized and the sequence analysis of these isolates exhibited 7 different species such as; *T. harzianum, T. viride, T. brevicompactum, T. ovalisporum, T. erinaceum, T. asperellum* and *T. orientalis*. Similar findings were supported by the investigators (39, 40). *Trichoderma* isolates of different bio-control capabilities were also characterized in three main clades with ITS sequence analysis (21). The genetic variability in sixty nine isolates of *Trichoderma*, and their phylogenetic analysis were grouped more than 50% of the potential *Trichoderma* strains with the sequence data from a fragment of the translation elongation factor-1 alpha gene (tef1) (41). The complementing findings were also supported by the scientist (42). However, the phylogenetic analysis in tef1 extended wider differentiation among the *Trichoderma* isolates within and among groups of closely related species. The higher level of variability was found in tef1 than in the rDNA of the ITS region as supported (43, 44, 45). *Trichoderma* isolates as *T. virens* (11 isolates), *T. asperellum* (15), *T. harzianum* (14) and *T. longibrachiatum* (32) were identified on the basis of the DNA sequence data analysis of the translation elongation factor 1-alpha gene (tef1) and ITS region (23). *Trichoderma* isolates were not fairly differentiated in ITS 1 and 2 regions analysis while these were clearly separated with tef1 sequences analysis into different clusters, while the sequence analyses with tef1 was tested as better marker to distinguish *Trichoderma* species because of greater transition ratio and evolutionary divergence.

Greater than 80% boot strap support values were remarked in identification of *Trichoderma* species with the corresponding representative strain in phylogenetic analysis based on tef1 (46). Well supported bootstrap value of 87% was grouped with the representative strain TR 48 of the new species *T. asperellum* (39). These observations were in consonance of the researchers (47, 48). It was necessarily important that *Trichoderma asperellum* were not found distinctly among 45 crop species of 49 crops. Even though the 105 locations were free from *Trichoderma* species. Consecutively, the rhizoshere soil did identify *Trichoderma asperellum* in 4 different locations throughout the country except the areas of saline belt (Khulna, Satkhira, Bagehat, Patuakhali, Pirozpur, Jhalokati, Barisal and Cox’s Bazar). This finding was supported by the researcher (49) who observed that spore density of *Trichoderma harzianum* was gradually reduced as the increasing of salinity and sporulation was very weak at 8 g L^-1^ of Nacl, while significant reduction of mycelium dry weight was also obtained at 8 g L^-1^. Therefore, the bioagent did survive in non-saline area of medium. Besides this, the *Trichoderma* isolates of BHUT6 and BHUT8 showed greatly salt tolerant in vitro condition and they were grown at 1400 mM NaCl concentration. Of these, the isolate of *Trichoderma* BHUT8 was determined as *Trichoderma asperellum* (accession number KU533735) (50) which were not recognized with the identified accession of *Trichoderma* isolate in our study. *Trichoderma* isolates were identified as *Trichoderma asperellum* of 4 crops in different locations. All the *Trichoderma* isolates resulted in growing and sporulating at different types of medium such as P^H^ value and temperatures in physiological features. Four isolates such as Sadar, Moulvibazar (Rice), Sadar, Mymensingh (Sweet gourd), Chapra, Chapai nawabganj (Sesmae) and Nayanpur, Lalmonirhat (Maize) were determined as *Trichoderma asperellum* in ITS region and *tef1* nucleotide sequencing. It was evident that the *Trichoderma* isolates were native to Bangladesh as *Trichoderma asperellum* strain in four crops (Rice, Sweet gourd, Sesame and Maize) that were found to be esteemed as higher potential in physiological and molecular characters.

## MATERIALS AND METHODS

### Experimental site

The experiments were conducted during January 2015 to December 2018 at the Eco-friendly Plant Disease Management Laboratory, Molecular Plant Pathology Laboratory and Bio-safety Laboratory of the Department of Plant Pathology, and Net house of Prof. Golam Ali Fakir Seed Pathology Center, Bangladesh Agricultural University, Mymensingh.

### Isolation, purification and preservation of *Trichoderma* of rhizosphere soil samples in 30 agro-ecological zones (AEZ) of Bangladesh

*Trichoderma* was isolated from soil samples using the soil dilution plate technique method (51). The surface of the working area was disinfected with cotton soaked in methylated spirits (70%). Hands were also disinfected using the same process and the glassware (test tubes, Petri dishes, pipettes, and beakers) were sterilized in a dry oven.

### Collection of soil samples

Rhizosphere soil samples were collected for isolation of *Trichoderma* species from 30 AEZs in 36 different districts (Satkhira, Khulna, Bagerhat, Jhenaidah, Jessore, Kushtia, Barisal, Patuakhali, Jhalokati, Pirojpur, Comilla, Chittagong, Cox’s Bazar, Khagrachari, Sylhet, Moulvibazar, Dhaka, Gazipur, Manikganj, Rajbari, Faridpur, Mymensingh, Tangail, Jamalpur, Rajshahi, Natore, Pabna, Sirajganj, Bogra, Chapainawabganj, Rangpur, Gaibandha, Kurigram, Lalmonirhat, Dinajpur and Thakurgaon) of Bangladesh. Altogether 109 soil samples were collected randomly from the rhizosphere of 49 crops (rice, spinach, brinjal, lentil, sundari, cane, mustard, marigold, tomato, bitter gourd, taro, red amaranth, banana, okra, jute, papaya, malta, tobacco, betel leaf, maize, nut grass, tea, asthma plant, litchi, bottle gourd, cotton, sissoo, onion, wheat, cabbage, hedge bean, broccoli, sweat gourd, german grass, napier grass, data shak, pointed gourd, pineapple, watermelon, sugarcane, jackfruit, sweat potato, dhaincha, sesame, garlic, chilli, mango, chickpea and natishak) in 109 locations (Satkhira (Debhata, Shyamnagar, Kaliganj, 3), Khulna (Rupsha), Bagerhat Sadar (3), Jhenidah (Kaliganj), Jashore (Jhikargacha), Kushtia (Mozompur), Barisal (Bakerganj, 2), Patuakhali (Mirzaganj, 2, Dashmina, 2), Jhalokati (Nalchity), Pirojpur (Kawkhali), Comilla (Daudkandi, 3, Chandina, 1), Chittagong (Lohagara, 3), Cox’s Bazar (Chakaria, 2), Khagrachari (Panchari, 2), Sylhet (Sadar, SAU gate hill, Haripur, Jaflang, Jaintapur, 4), Moulvibazar (Sadar, 2, Kulaura, 1, Barlekha, 2), Dhaka (Savar, 2, Banani, 1), Gazipur (Sadar, Sreepur, Rajendrapur, 3), Manikganj (Paturia), Rajbari (Goalanda), Faridpur (Bhanga), Mymensingh (Sadar, BINA field (5), Sadar, Hort. Field, BAU (9), Sadar, Dairy farm, BAU (2), Sadar, Sutia khali, BAU (3), Sadar, Sutia khali, BAU (2)), Tangail (Madhupur, Sadar, 2), Jamalpur (Sadar, 2), Rajshahi (Sadar, BARI centre, 2), Natore (Sadar, Baraigram, 2), Pabna (Ishwardi, 3), Sirajganj (Sadar), Bogra (Shibganj, 5), Chapainawabganj (Sadar, 3), Rangpur ((Sadar, 3, Mithapukur, 3), Gaibandha (Gobindaganj), Kurigram (Sadar, 5), LalmonirHat (Sadar, 3), Dinajpur (Sadar, 2) and Thakurgaon (Sadar, Pirganj, 2) of Bangladesh. The soil samples were kept in a refrigerator (4 °C) in the laboratory until use.

### Preparation of soil dilution

Soil (1 g) was kept in a test tube with 9 ml of sterile water and stirred thoroughly for a few minutes to obtain a uniform 10 ° dilute soil suspension. It was used as stock suspension. The stock suspension (1 ml) was transferred with a sterile pipette into a second test tube in 9 ml sterile water and it was shaken thoroughly, while a 10^−1^ dilute soil suspension was obtained. The same process was repeated for further dilution as 10^−3^.

### Isolation of *Trichoderma* from soil and preparation of pure culture

Twenty ml of warm melted PDA medium was (approx. 45 °C) poured in each sterile One ml of diluted soil sample (10^−3^) was kept at the center of a plate on PDA and it was spreaded with a glass rod. Each was inoculated with 1 ml of diluted sample and repeated with every soil sample. The inoculated PDA plates were incubated for 7-10 days at room temperature (25 ± 1 °C). Plates were observed for *Trichoderma* colonies after incubation of 3 days. The growing margin of the *Trichoderma* colony was cut into 5 mm discs with a cork borer. The discs were carefully kept in a new PDA plates to produce a pure culture of *Trichoderma* and the new plates were also incubated. The pure culture of *Trichoderma* was found to grow sufficiently within 3 days of incubation. The cultures were prepared as sub-culture to PDA plate and were transferred to PDA slants for preservation as separate culture for each. The complete grown of *Trichoderma* in slants was preserved in the refrigerator at 4 °C for further use. The isolates were identified following the key (52).

### Morphological and physiological characterization of collected *Trichoderma*

The isolates were characterized morphologically and were purified through mycelia block culture, and were maintained on acidified potato dextrose agar (APDA) medium at 28 ± 1 ^0^C (incubation) for studying cultural characteristics (53). Radial mycelia growth of the isolates was studied following the method (54). Colony diameter was measured up to 6 days after inoculation of 24 hours. The colonies were filled the plant and sporulation was occurred within 6 days. After 24, 48 and 72 hrs of inoculation, the radial mycelia growth was measured as the mean of two perpendicular diameters and mean of three replications was taken as growth of each isolate.

The number of colonies was recorded per gram of soil. The colonies were regular or irregular in shape. Colony colors, black/ white/ grey/ blackish/ whitish/ blackish white/ whitish black were recorded. After 7 days of incubation of a single culture on PDA mean radial growth of the isolate was measured in the method (55).

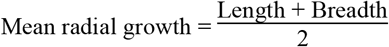

**C**ompact or loose characters were determined on the basis of compactness of the colony. The surface of the culture was recorded as smooth/glistening/rough/wrinkled/ dull etc.

The opacity of the culture such as transparent (clear), opaque, translucent (like looking through frosted glass) etc. were determined.

#### Number of spores per ml

Conidia per ml after 7, 14 and 21 days of incubation were determined following the formula (54).]

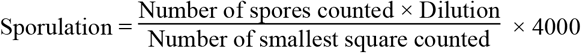

### Preparation of inocula of *Trichoderma* isolates, selection of isolates and measurement of spore density

Eighty eight *Trichoderma* isolates were collected from rhizosphere soil of 49 crops in109 locations of the country. The isolates were categorized as fast, medium and slow growing on the basis of their growth rate. Fifteen isolates were selected for further study as fast growing among 88 isolates. The selected *Trichoderma* isolates were grown on PDA medium as described (57). The mycelial blocks (1 cm^2^) of pure cultures were transferred into the medium. The plates were incubated at 30 °C for growth and sporulation.

### Measurement of spore density

The spore density was determined using of 100 ml water in a test tube and about 10 ml was poured in each PDA plate with 7, 14 and 21 days old culture of *Trichoderma*. The colony was scraped smoothly and gently with a scalpel or slide to collect conidia. The suspension was taken into a beaker from the medium and stirring was continued with a glass rod. It was repeated few times. The volume of the beaker with conidial suspension was made 100 ml with sterile water and 1 drop of Tween-20 was added to it and stirred to disperse well. One drop of suspension of the solution was taken on the centre of haemocytometer and a cover slip was kept on it. The spores/ml was counted under microscopic power of 40X with haemocytometer using the block system.

### Physiological characterization of *Trichoderma*

The isolates of *Trichoderma spp*. were characterized physiologically. The isolates of *Trichoderma* were purified through mycelia block culture and were incubated on Acidified Potato Dextrose Agar (APDA) medium at 30 ^0^C in studying cultural characteristics (53).

### Effects of temperature on mycelial growth of *Trichoderma spp*

Seven levels of temperature viz., 4 ^0^C, 15 ^0^C, 20 °C, 25 °C, 30 °C, 35 °C and 40 °C were tested to find out the effect of temperature on mycelial growth of *Trichoderma spp*. The experiment was conducted in three incubators (Froilabo, model no. 69330 MEYZIEU-FRANCE) having each with different temperature. The influence of temperature levels on mycelial growth was identified on the basis of colony diameter of mycelium. The fungus was grown on PDA plates for radial colony growth. Forty five petri dishes of solidified PDA were inoculated with the inocula of the fungus following the procedure (53). After inoculation, the petri dishes were kept in the incubators for 7days.

### Effects of pH on mycelial growth of *Trichoderma*

Four levels of p^H^ at 6.0, 7.0, 8.0 and 8.5 were tested. PDA was prepared and three PDA plates were used for each treatment. Before cooking, all ingredients were thoroughly mixed in water on a magnetic stirrer. The p^H^ of the medium was adjusted to required level with an electrode p^H^ meter. NaOH of 0.1N and 0.1N HCl were added to increase and decrease p^H^ levels, respectively. The p^H^ level was adjusted at 6.5 after extracting of potato, while the medium was boiled on a microwave oven for melting the agar powder. This media was used as controlled condition. The PDA plates were inoculated with mycelial disks of the fungus. The inoculated plates were incubated at 30 °C. Data on radial colony diameter was recorded.

### Statistical analysis

The recorded data on different parameters were analyzed statistically using Web Agri Stat Package computer program to find out the significant variation of experimental treatments. The difference between the treatments means were evaluated with Duncan’s Multiple Range Test (DMRT) following the procedure (58).

### Molecular characterization of selected *Trichoderma*

Four isolates of the 15 *Trichoderma* isolates were selected on the basis of their better spore density and tolerable in high temperature in vitro condition. These were characterized using molecular marker of ITS4, ITS5 and *tef1, tef2*.

### Preparation of broth culture to harvest mycelia of *Trichoderma spp*

*Trichoderma* isolates were cultured individually in 150 ml conical flasks with 100 ml liquid potato dextrose culture medium for DNA extraction. Hypha were collected from potato dextrose broth (PDB) with mycelia and incubated for 7 days in the incubator at 25 ± 1 ^0^C on filter paper in a Buchner funnel, and it was washed with distilled water, frozen, and lyophilized. After harvest, mycelia were wrapped with aluminum foil sheet for each isolate separately and kept at 4 ^0^C until prior to the genomic DNA isolation.

### Genomic DNA extraction

The genomic DNA of each *Trichoderma* isolate was extracted from harvested mycelia of 3 days old culture following the protocol of Wizard ^®^ Genomic DNA Purification Kit (Year 2018). Forty mg of mycelia was cut into small pieces and kept in 1.5 ml eppendorf tube and liquid nitrogen was added to grind with micro pestle. Six hundred µl of Nuclei Lysis solution was mixed and vortexed for 20 seconds for proper mixing and incubated at 65 °C for 15 minutes in hot water bath for digestion. The solution of 3µl of RNase was added and incubated at 37°C for 15 minutes. Then the sample was cooled at room temperature for 5 minutes. Protein precipitation solution of 200 µl was mixed and vortexed gently, and centrifuged at 15,000 rpm for 3 minutes. The supernatant was transferred in clean tube without disturbing the lower portion and 600 µl was put together in room temperature. Then isopropanol was added to the supernatant and shaken slowly. Mixed by inversion was done and centrifuged at 15,000 rpm for 3 minutes to produce precipitation of the cell debris. Six hundred µl of 70% ethanol was mixed in blended supernatant at room temperature and centrifuged at 15,000 rpm for 2 minutes. The pellet was air dried in order to evaporate the ethanol. DNA rehydration solution of 25µl was added and mixed gently by finger tapping. Finally the DNA solutions were preserved overnight at 4 °C.

### Determination of DNA concentration

The spectrophotometer was set at 260 nm for quantification of DNA. A square cuvette (the zero or blank cuvette) was filled with 2 ml double distilled water and kept in the cuvette chamber. Then the absorbance reading was adjusted to zero for standardization. The test samples were prepared with 2 μl of each DNA sample in the cuvette of 2 ml sterile distilled water and mixed comprehensively by pipetting. The absorbance reading was taken at 260 nm and the cuvette was rinsed with sterile water. The reading for each sample was recorded in the same way. The original concentration was determined by using the above absorbance readings in the following formula:

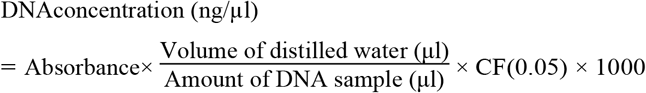

### PCR amplification of the ITS region (ITS4 and ITS5) and *Tef* region (*Tef*1-728F and *Tef2*-986R)

Specific primers such as ITS4, ITS5, *tef*1-986R and *tef*2-728F were applied in matching the template DNA of the *Trichoderma* to confirm the *Trichoderma* through PCR. PCR reactions were determined on each amplified DNA with the sequence of TCCTCCGCTTATTGATATGC (5’-3’) and GGAAGTAAAAGTCGTAACAAGG (5’-3’) both with 300-650 bp in the primer of ITS4 and ITS5, respectively (53), while the sequence CATCGAGAAGTTCGAGAAGG (5’-3’) and TACTTGAAGGAACCCTTACC (5’-3’) were found in the primer of *Tef*1-728F and *Tef2*-986R both with 250-350 bp. The PCR reactions were performed in 25 µl of reaction mixture for each DNA template which composed of GoTaqR G2 green master mix of Taq polymerase buffer and dNTPs, forward primer 1.0 µl, nucleus free water 9.50 µl and DNA template 1.0 µl. The amplification was carried out using of T 100 Thermal Cycler (Bio-Rad, Hercules, CA, USA) following the profile (39). The thermal cycling profile was started at 95 °C for 5 minutes in pre-denaturation followed by 35 cycles at 94 °C for 1.5 minute, 2 min annealing at 55 °C, extension at 72 °C for 3 minutes and 5 minutes for 72 °C in final extension for all amplified fragments in ITS region. Thermal cycling profile of TEF region was begun with 2 minutes for 94 °C in pre-denaturation followed by 30 cycles at 98 °C for 10 seconds in denaturation, 30 second annealing at 55 °C, extension for 35 second at 72 °C and 10 minutes for 72 °C in final extension of all amplified fragments. Amplified DNA of all isolates and total 7 µl of 100 bp DNA ladder were used in the gel electrophoresis at 80 Volt for 55 minutes. The gel was stained with ethidium bromide for 30 minutes at room temperature and it was transferred from the Ethidium bromide tray and kept on the UV transilluminator for image documentation of DNA bands and photographed with the Gel Documentation System.

### DNA sequencing and analyses of nucleotide sequences

DNA sequencing was done following the standard protocols (dna.macrogen.com) for the ABI 3730×1 DNA genetic analyser (Applied Biosystems, Foster City, CA, USA) with BigDye ® Terminator v1.1 and 3.1 Cycle Sequencing Kits. The necleotide sequences were analyzed with online bioinformatics tools. The DNA sequences were compared with other *Trichoderma spp*. available in the NCBI database of Basic Local Alignment Search Tool (BLAST) algorithm to identify closely related sequences (http://blast.ncbi.nlm.nih.gov/Blast.cgi).

### Phylogenetic analysis

Phylogenetic analysis was conducted following the methods of MEGA7 Version 5.22 [4].

## ACKNOWLEDGEMENTS

This research work was supported by the HEQEP-AIF funded project entitled “Strengthening Postgraduate Research in Plant Protection for Sustainable Crop Production”, Department of Plant Pathology, Bangladesh Agricultural University, Mymensing-2202, Bangladesh.

